# The mood stabilizer lithium slows down synaptic vesicle cycling at glutamatergic synapses

**DOI:** 10.1101/780866

**Authors:** Willcyn Tang, Bradley Cory, Kah Leong Lim, Marc Fivaz

**Affiliations:** National Neuroscience Institute, 11 Jalan Tan Tock Seng, Singapore 308433; Stem Cell & Gene Editing Laboratory, University of Greenwich, Faculty of Science and Engineering, Kent ME4 4TB, UK

**Keywords:** bipolar disorder, synaptic vesicle cycle, glutamatergic synapse, neurotransmission

## Abstract

Lithium is a mood stabilizer broadly used to prevent and treat symptoms of mania and depression in people with bipolar disorder (BD). Little is known, however, about its mode of action. Here, we analyzed the impact of lithium on synaptic vesicle (SV) cycling at presynaptic terminals releasing glutamate, a neurotransmitter previously implicated in BD and other neuropsychiatric conditions. We used the pHluorin-based synaptic tracer vGpH and a fully automated image processing pipeline to quantify the effect of lithium on both SV exocytosis and endocytosis in hippocampal neurons. We found that lithium selectively reduces SV exocytic rates during electrical stimulation, and markedly slows down SV recycling post-stimulation. Analysis of single bouton responses revealed the existence of functionally distinct excitatory synapses with varying sensitivity to lithium ― some terminals show responses similar to untreated cells, while others are markedly impaired in their ability to recycle SVs. While the cause of this heterogeneity is unclear, these data indicate that lithium interacts with the SV machinery and influences glutamate release in a large fraction of excitatory synapses. Together, our findings show that lithium down modulates SV cycling, an effect consistent with clinical reports indicating hyperactivation of glutamate neurotransmission in BD.

## Introduction

Bipolar disorder (BD) is a common psychiatric illness characterized by prominent mood swings with episodes of mania and depression. BD is the 6th leading cause of disability among adolescents and young adults (1) ― without treatment, 15% of BD patients commit suicide (2). Since its introduction by John Cade in 1949 (3), lithium has remained the single most effective treatment for BD (4). The discovery of lithium’s effect on mania was purely accidental. Based on the misconception that there was some connection between mania and urea, John Cade injected uric acid into guinea pigs and used lithium urate as a solubilization agent. To his surprise, this produced a calming effect, instead of the predicted excitement. Through a series of carefully controlled experiments, Cade was able to identify the lithium salt as the active compound. He administered lithium to 10 manic patients who responded remarkably well (3), a breakthrough that signaled the discovery of the first effective drug to treat mental illness. Meta-analysis of randomized controlled trials have since shown that lithium is effective against both the manic and depressive phases of BP ― lithium reduces the risk of manic relapses by 38% and depressive relapses by 28% (5). The drug also decreases the risk of suicide in patients with mood disorders by more than 50% (6). However, only a fraction of BP patients respond to lithium and the drug suffers from low therapeutic index and adverse side effects (weight gain, possible renal dysfunction), creating an unmet need for better mood stabilizers. Moreover, despite the use of lithium in the clinic for more than 50 years, its mechanism of action is poorly understood. Several molecular targets have been identified ― Gsk3 (7), inositol phosphatase (8), CRMP2 (9), TrkB (10), and Protein kinase C (11) ― but it is not clear how these proteins and pathways contribute to BD pathogenesis. The difficulty in assigning a mechanism of action for lithium is linked to our poor understanding of BD’s etiology. Increasing evidence, however, points to glutamate hyperexcitability as a central feature of this mental condition. Functional imaging studies show widespread increase in combined glutamate and glutamine levels in the brain of BD patients (12,13) and elevated glutamate levels have also been reported in prefrontal cortex of BD postmortem brains (14). Perturbed glutamate neurotransmission in BD is further supported by several imaging and postmortem studies describing altered levels of postsynaptic glutamate receptors. Notably, induced pluripotent stem cells (iPSCs)-derived hippocampal neurons from BD patients are hyperexcitable (15), a phenotype which is selectively reversed by lithium in neurons from patients that respond to lithium therapy (15). Consistent with a role in reducing glutamatergic neurotransmission, lithium protects against excitotoxicity by inhibiting NMDA receptors (16) and its antimanic effect is mediated by BDNF/TrkB-dependent synaptic removal of AMPA receptors, at least in a mouse model of hyperactivity (10). While several studies have demonstrated a role of lithium in dampening the activity of postsynaptic glutamate receptors, comparatively little is known about the effect of lithium on presynaptic function. Here, we investigate the impact of lithium on synaptic vesicle (SV) exo- and endocytosis at individual presynaptic terminals of hippocampal neurons using an imaging assay based on pHluorin (17) fused to the glutamate transporter VGlut1 (vGpH) (18). We show that lithium slows down SV exocytosis, but does not alter rates of SV endocytosis during electrical stimulation. Lithium decreases, however, rates of SV reuptake post-stimulation suggesting distinct modalities of SV endocytosis during and after action potential (AP) firing. Collectively, our findings show that lithium alters the kinetics of SV cycling at excitatory synapses, an effect expected to decrease glutamatergic neurotransmission, particularly during bursts of high frequency firing. These results identify the SV cycling machinery as a functional target of lithium and highlight a novel mechanism by which lithium may counterbalance elevated glutamate neurotransmission in BD patients.

## Results

We used the synaptic reporter vGpH (18) to examine the effect of lithium on presynaptic function in hippocampal neurons. vGpH consists of the super-ecliptic pHluorin (17) fused into the first lumenal loop of the VGlut1 transporter (18,19). In this configuration, pHluorin faces the acidic lumen of SVs and its fluorescence is quenched in resting neurons. During action potentials (APs), exocytosis of SVs exposes pHluorin to the neutral pH of the extracellular environment leading to a ∼ 20 fold increase in fluorescence intensity (20). Following endocytosis and re-acidification of SVs the fluorescence of vGpH returns to baseline. Because re-acidification of SVs is not rate-limiting (20) changes in vGpH intensity during AP firing directly reflect the balance between SV exo- and endocytosis.

We have previously developed an image processing pipeline to automatically identify functional (responsive) synaptic boutons and compute their SV exo- and endocytic rates from vGpH time series (19). Isolation of responsive boutons is achieved by segmentation of a differential vGpH image that selectively displays vGpH intensity increase during stimulation (Figure 1a-c). Hippocampal neurons were stimulated with two consecutive trains of action potentials (APs) 5 min apart, to allow full regeneration of the SV releasable pool (Figure 1d). Each train consists of 300 APs delivered at 10Hz (36.5°C), under conditions that do not deplete the SV pool or saturate the SV endocytic machinery (21). The shape of the first vGpH transient is governed by relative rates of SV exocytosis and reuptake. The second AP train is delivered in the presence of the vATPase blocker bafilomycin (baf) that prevents re-acidification of SVs and thus allows selective imaging of SV exocytosis (19,22). The endocytic component is then computed by subtracting the first vGpH transient from the vGpH response after baf treatment (Figure 1d,e). Intensity traces show substantial synapse-to-synapse variability within a single field of view (FOV), even after normalization of vGpH expression using NH4Cl (Figure 1d), a behavior consistent with previous reports (19,22,23).

**Figure 1.**
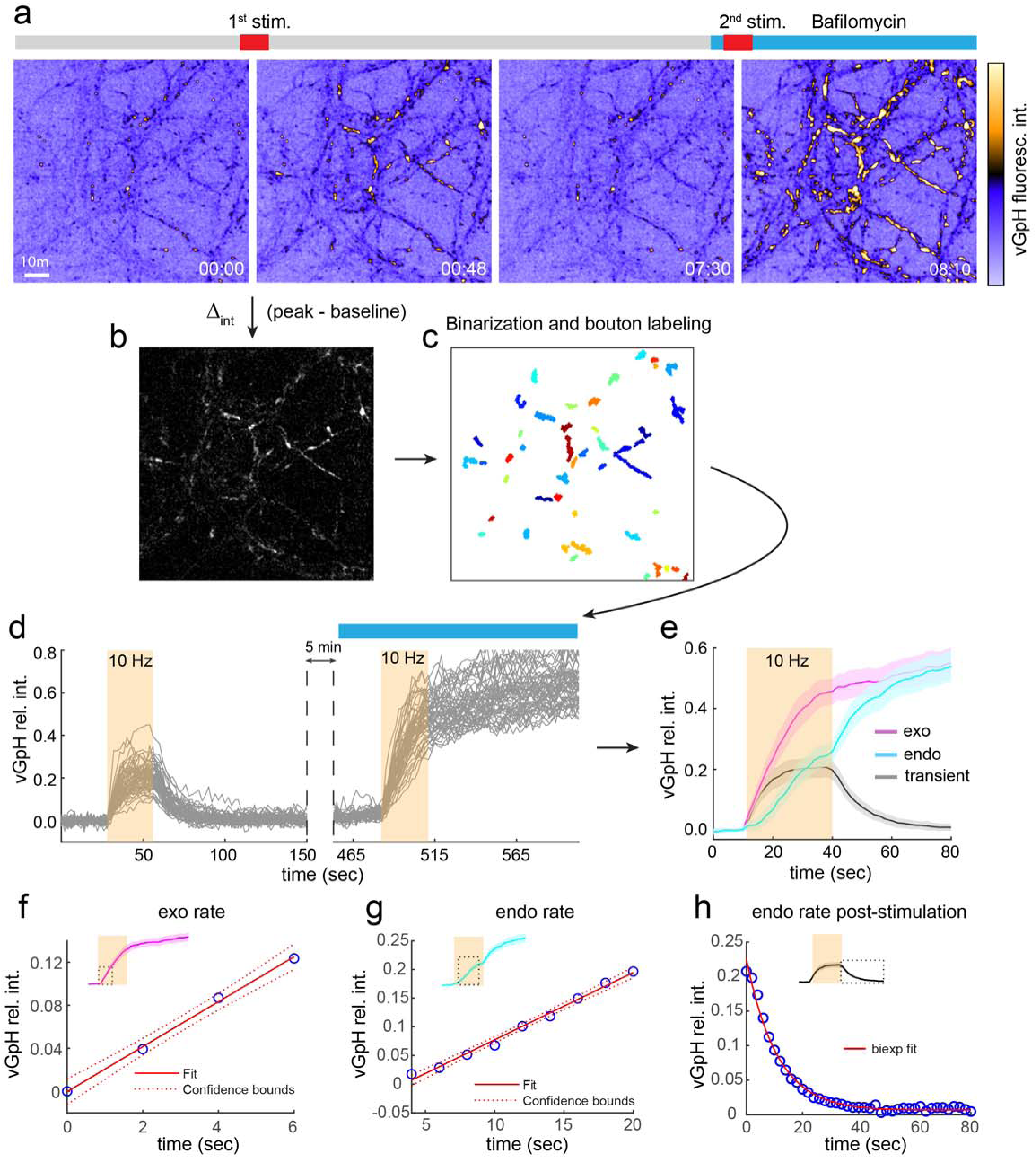
Image processing pipeline. (a) Snapshots of a vGpH time series showing the first and second vGpH response before and after baf treatment. Neurons were stimulated with two consecutive trains of 300 APs at 10Hz. Time is in min and sec. (b,c) Segmentation of responsive boutons based on the difference of vGpH intensity before and at the peak of the response during stimulation. (d) vGpH traces for each synaptic bouton detected in the field. Traces were normalized for vGpH expression by NH4Cl at the end of the time series (not shown). (e) The first vGpH transient overlaid with the mean SV exocytic and endocytic components computed from traces shown in (d). Shaded error bar indicates 95% CI (confidence interval). (f-h) Estimation of SV exocytic (f) and endocytic rates (g), and endocytic time constant post-stimulation (h).

We measured SV rates from an average vGpH trace generated for each FOV. Six to thirteen FOVs were analyzed for each treatment and three SV cycling parameters were measured: SV exo and endo rates during stimulation and time constant for SV endocytosis post-stimulation (Figure 1f-h). We used different time windows to fit the exo- and endo traces during AP firing based on the shape of these traces (Figure 1f,g) so exo and endo rates are not directly comparable to each other.

Neurons incubated with 5mM LiCl for 24h ― a concentration within the therapeutic range (24) ― showed a 28% decrease in SV exo rates (p = 0.036, Mann-Whitney U test) compared to control cells (Figure 2a). Lithium has no apparent effect of SV endo rates during stimulation (Figure 2b), but appears to slow down SV uptake post-stimulation as reflected by an increase in the endocytic time constant (Figure 2c, p = 0.13, Mann-Whitney U test), although the p-value is above the 5% cutoff after correction for multiple hypothesis testing (see methods). Longer (48h-72h) exposure to lithium exacerbated these effects, with a 31% reduction in SV exo rates (Figure 2d, p = 0.001, Mann-Whitney U test) and a 32% increase in the endocytic time constant post-stimulation (Figure 2f, p = 0.018, Mann-Whitney U test), while SV endo rates during stimulation remained unperturbed (Figure 2e).

**Figure 2.**
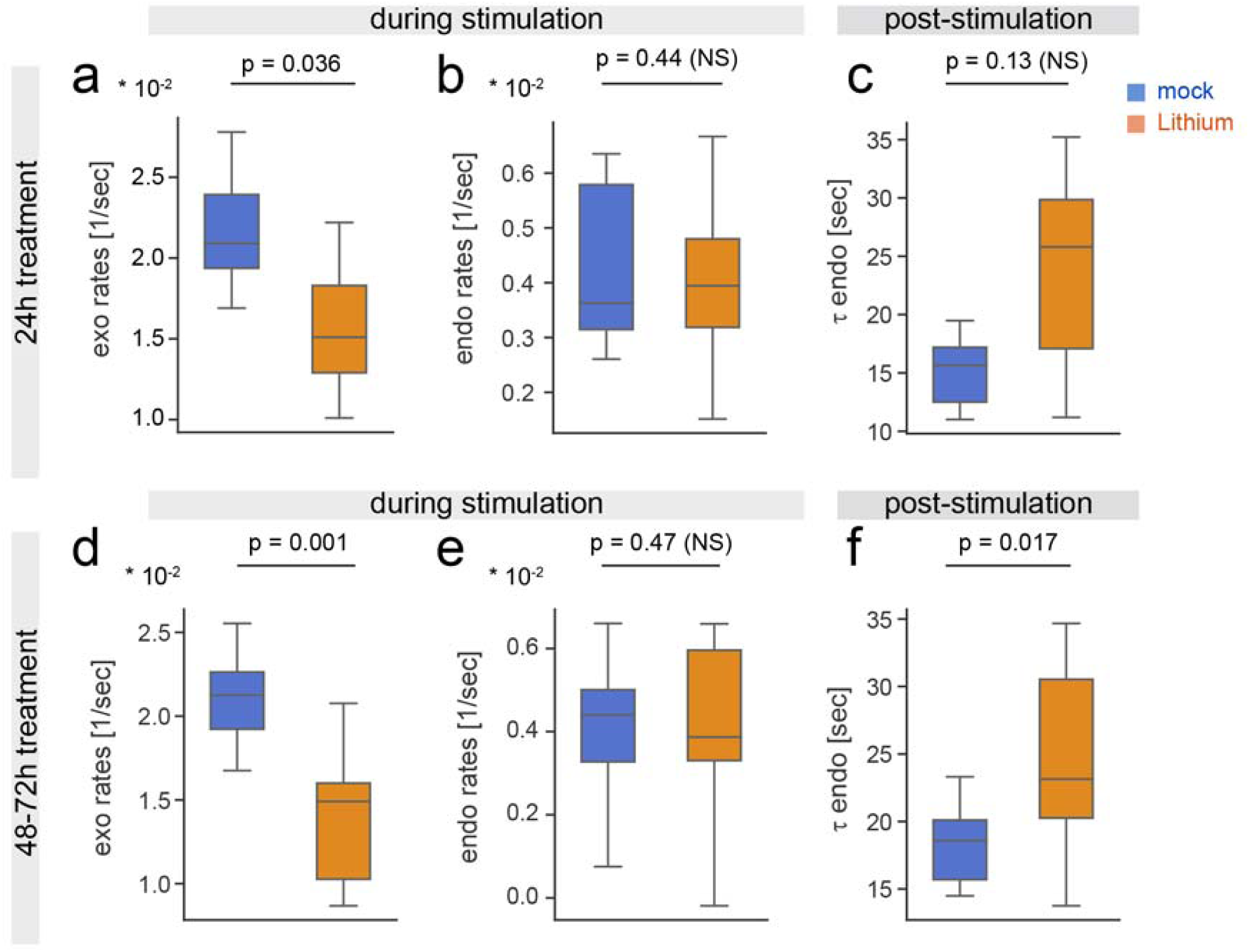
SV cycling parameters in control and lithium-treated neurons. Boxplots showing SV exo and endo rates and endo time constants post-stimulation in control and lithium-treated neurons 24h (a-c) and 48-72h (d-f) after treatment. Each dataset consists of 6-13 fields of view (more than 1000 boutons each) originating from at least 4 independent experiments. P values were obtained by a Mann-Whitney U test and corrected post-hoc for multiple hypothesis testing.

The effect of lithium on post-stimulation SV endocytosis, but not on SV uptake during stimulation is puzzling. One possible explanation is that the endocytic phenotype during stimulation is masked by a particularly large variability in endocytic rates across experiments (Figure 2b,e). Alternatively, SV endocytosis during and after stimulation could operate under different regimes (see discussion) and could, therefore, be differentially affected by lithium. To test these two scenarios, we analyzed SV cycling in individual FOVs to eliminate field-to-field variability. The vGpH endocytic trace shows a clear biphasic behavior with a boost in SV endo rates immediately after stimulation (Figure 3a, see also Figure 1e). Such a biphasic behavior is consistent with two modes of SV endocytosis ― during and post-stimulation ― and is largely attenuated in lithium-treated neurons (Figure 3d). The endocytic component in these cells increases linearly with no abrupt change in rate during and after stimulation (Figure 3d). Accordingly, vGpH decay in control cells (after stimulation) is best fit with a biexponential function with fast and slow time constants (Figure 3b,c), while vGpH decrease in lithium-treated neurons resembles more a monoexponential decay (Figure 3e,f). Comparison of time constants shows that the initial (fast) rate of SV reuptake (the one that dominates the endocytic response) is slower in lithium-treated cells (Figure 3g), in agreement with the aggregate data (Figure 2c,f). Endocytic rates during stimulation, however, are slightly higher in lithium-treated cells (Figure 3g). Finally, exocytic rates are severely reduced by lithium (Figure 3g), consistent with the aggregate data (Figure 2a,d). Together, these data indicate that lithium can specifically slow down SV reuptake post-stimulation without decreasing SV endocytosis during APs. Lithium thus selectively impairs SV exocytosis during AP firing causing an imbalance between SV exo- and endocytosis (Figure 3h).

**Figure 3.**
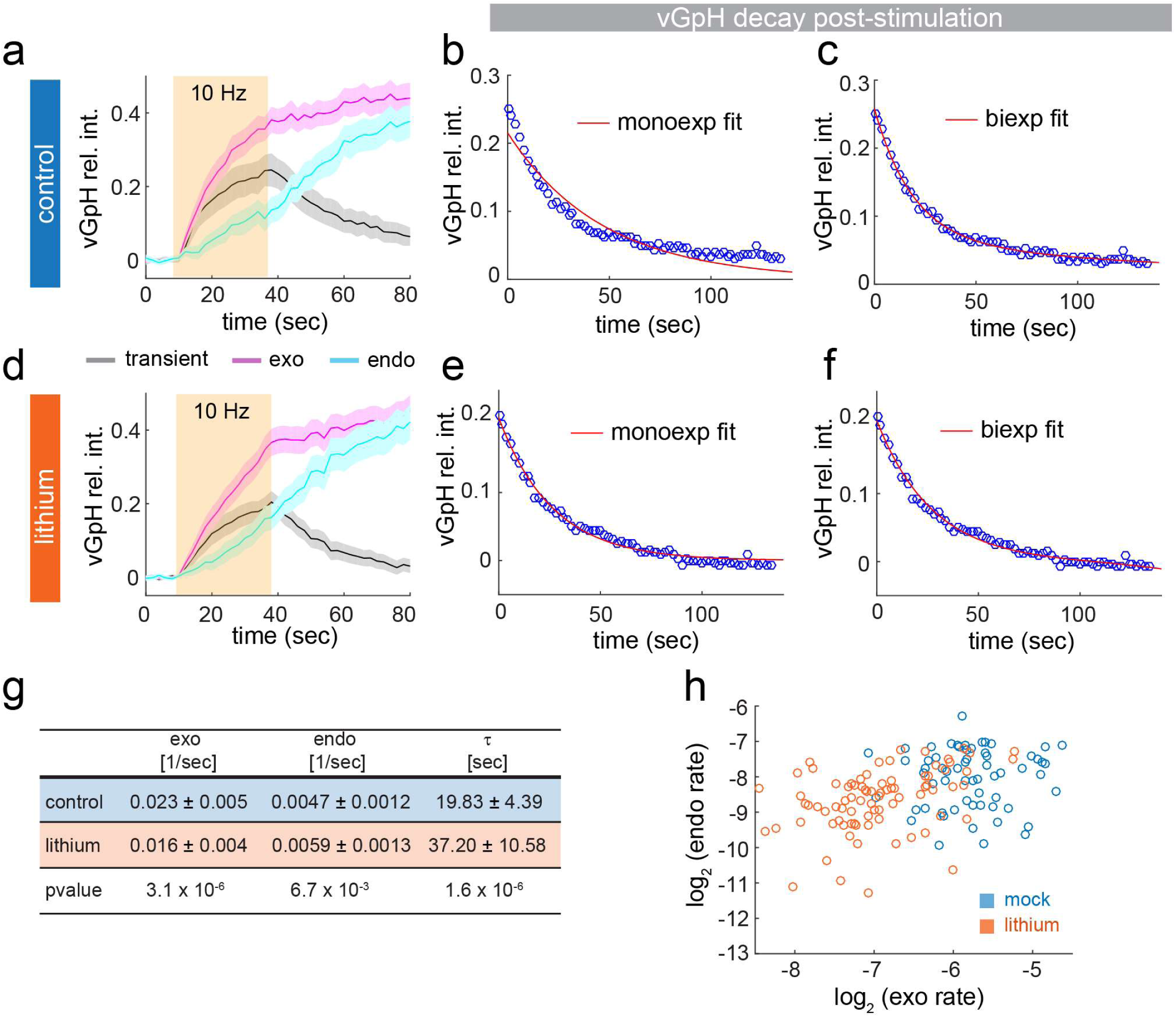
Lithium differentially affects SV endocytosis during and after stimulation. Analysis of SV cycling in a single control FOV (a-c, g) and lithium-treated FOV (d-f, g). SV endocytosis post-stimulation is best fit with a biexponential function (b,c). In lithium-treated neurons, SV endocytosis post-stimulation is well fit with a monoexponential function (e). A bi-exponential fit (f) leads to two components with similar time constants (not shown), reflecting the absence of a fast endocytic component (see text). (g) Mean SV cycling parameters with 95% CI. Only fast time constants are shown. P values were obtained by a Mann-Whitney U test and corrected post-hoc for multiple hypothesis testing. (h) Scatter plot showing the typical relationship between exo and endo rates for each synaptic bouton in a different pair of control and lithium-treated FOVs. As in (a-g), lithium has little impact on endocytic rates, but markedly decreases exocytic rates during stimulation, causing a severe imbalance in SV cycling.

We next examined the impact of lithium on SV cycling at individual boutons. We first compared vGpH time series at single presynaptic terminals in both control and lithium-treated neurons using a heatmap display (Figure 4a). Traces for each group originate from a single FOV comprised of roughly the same number of boutons. While vGpH transients appeared relatively homogenous in untreated neurons, they exhibited marked differences in lithium-treated cells (Figure 4a). Many terminals failed to recycle SVs efficiently as reflected by protracted vGpH transients. Others showed vGpH responses indistinguishable from untreated boutons. To further compare vGpH transients in these two groups, we merged them into a single dataset and analyzed vGpH traces by unsupervised hierarchical clustering (Figure 4b). The algorithm identified two main clusters corresponding to lithium-treated (orange) and control (blue) boutons, with a small fraction (< 5%) of control boutons behaving as lithium-treated ones and a larger fraction (∼ 20%) of lithium-treated boutons behaving as control ones. Clustering in these two main groups correlates to a large extent with the duration of the vGpH transient. To specifically determine the effect of lithium on SV endocytosis post-stimulation, we plotted the distribution of time constants for these two groups. This probability analysis shows a narrow distribution of time constants for control cells, and a broader distribution shifted towards higher time constants (i.e. slower SV uptake post-stimulation) for lithium-treated cells (Figure 4c). A fraction of boutons (about 20%) displays time constants overlapping with those of control terminals (Figure 4c). We carried out this distribution analysis on multiple pairs of control and lithium-treated fields acquired on different days from independent neuron preparations. In each case, lithium slowed down SV reuptake post-stimulation (Figure 4c-f), although the fraction of boutons showing impaired endocytosis significantly varied across experiments ― some lithium-treated fields displayed a clear biphasic distribution of time constants with a large fraction of boutons (up to half) showing control-like endocytic responses (Figure 4d,f), while in others time constants are markedly higher, with little overlap between the two distributions (Figure 4c,e). Together these results indicate that glutamatergic terminals show varying degrees of sensitivity to lithium, with, at one extreme, boutons that are severely impaired in their ability to recycle SVs, and at the other end of the distribution, terminals that are insensitive to lithium.

**Figure 4.**
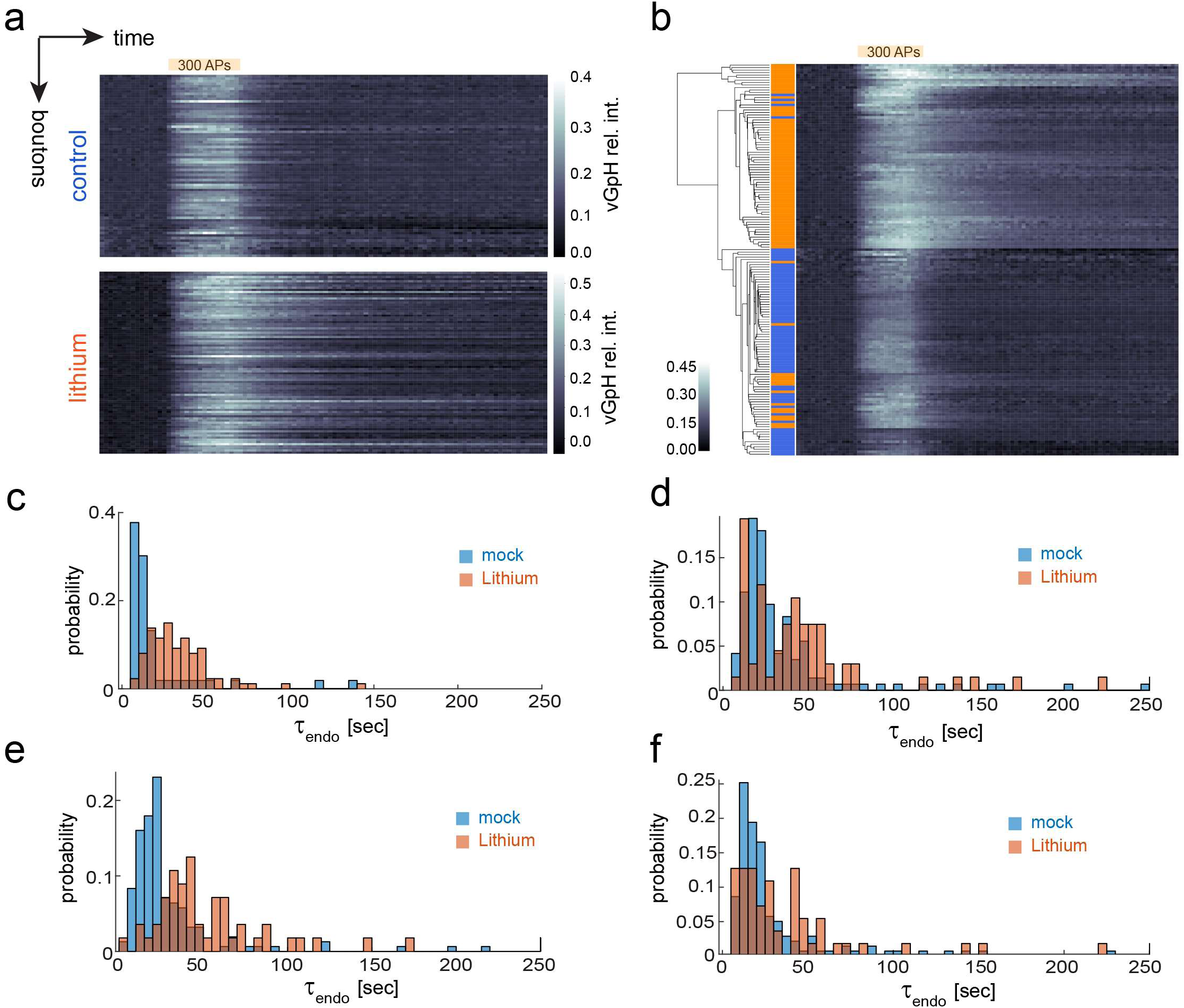
Identification of subtypes of excitatory synapses with varying sensitivity to lithium. (a) Heatmaps showing vGpH transients in individual boutons (rows) in control and lithium-treated fields. (b) Hierarchical clustering of vGpH transients shown in (a). Orange and blue rows correspond to lithium-treated and control boutons respectively. (c-f) Normalized distribution of endo time constants in different pairs of control and lithium-treated fields. (c) was generated from FOVs shown in (a,b).

## Discussion

Here we show that lithium decreases kinetics of SV exocytosis and endocytosis at hippocampal synapses with a specific effect on SV retrieval after, but not during AP firing. Our findings further suggest that SV endocytosis operates under different regimes during and after stimulation. Our evidence for distinct modes of SV endocytosis is based on bi-phasic kinetics of SV re-uptake during and after stimulation and on differential sensitivity of these two endocytic phases to lithium. In control (untreated) synaptic terminals we observed a notable increase in SV endocytosis immediately after stimulation. This acceleration is characterized by a biexponential decay in vGpH fluorescence with a fast time constant of ∼ 20 sec, in good agreement with previous measurements of SV endocytosis (25–27). Lithium abrogates this biphasic behaviour by suppressing this acceleration and leads to a slower monoexponential decay of vGpH. Such bi-phasic kinetics of SV endocytosis have not been directly observed before, perhaps because few studies monitor SV reuptake during and after AP firing. However, distinct modes of SV retrieval depending on the nature of the electrical stimulus have been previously described (18,26). For example, the Ryan lab has recently shown that SV endocytosis accelerates during short bursts of APs (up to 25), but slows down as the bursts lengthen (26). Acceleration is most prominent at 37°C and requires dephosphorylation of dynamin-1 by the calcium-dependent phosphatase calcineurin. Deceleration, on the other hand, appears to result from sustained elevation of Ca^2+^ levels in response to longer stimuli (26), and is consistent with earlier reports indicating an inhibitory function of Ca^2+^ in SV endocytosis (28,29). In our work, SV release was evoked by trains of 300 APs, at a frequency (10 Hz) that results in a pronounced rise of Ca^2+^ in the presynaptic compartment (19). It is therefore likely that SV endocytosis is slowed down under these conditions and accelerates again post-stimulation, as Ca^2+^ concentration decreases in the terminal.

What is the mechanism by which lithium slows down SV endocytosis post-stimulation? Two mechanisms of SV retrieval co-exist at nerve terminals; clathrin-mediated endocytosis (CME) which is dominant during low-intensity stimulation (25), and activity-dependent bulk endocytosis (ADBE), which is predominantly engaged during high-intensity stimulation (30). Interestingly, ADBE is regulated by Gsk3β, a serine/threonine kinase previously implicated in BD and a major target of lithium (7). A group of endocytosis proteins (the dephosphins) are dephosphorylated by calcineurin during stimulation and need to be rephosphorylated after stimulation for efficient recycling of SVs. One of these proteins, dynamin-1, is rephosphorylated by Gsk3β and its priming kinase cdk5 (31), a phosphorylation event which is both necessary and sufficient for ADBE. Because lithium inhibits Gsk3β, blockage of ADBE could be one mechanism by which this mood stabilizer slows down SV endocytosis post-stimulation.

In addition to its role in SV retrieval, lithium also decreases rates of SV exocytosis by ∼ 30%. How lithium impacts SV release remains unknown, but we can exclude an effect of the drug on the SV releasable pool since the vGpH signal reached the same plateau after bafilomycin treatment in control and lithium-treated cells, albeit with different kinetics (Figure 3a,d). Given the strong dependence of SV exocytosis on Ca^2+^ concentration, it is possible that lithium reduces Ca^2+^ influx and/or increases buffering of intracellular Ca^2+^ in the terminal. The N-type voltage operated Ca^2+^ channel Cav2.2 is the major source of AP-induced Ca^2+^ influx in presynaptic terminals of hippocampal neurons (19,23). Of note, Cav2.2 current density is enhanced by CRMP-2 (32), a phosphoprotein regulated by Gsk3β (33) and abnormally phosphorylated in iPSC-derived neurons from lithium-responsive BD patients (9). Thus, lithium could function through CRMP-2 to reduce Cav2.2-mediated Ca^2+^ entry and, consequently, slow down SV exocytosis.

This study further highlights the existence of functionally distinct types of synaptic terminals based on their response to lithium. We classified synaptic boutons based on the effect of lithium on SV endocytosis post-stimulation. The fraction of lithium non-responders is variable (20% to 50%) and is reminiscent of BD patients that don’t respond to lithium treatment. The mechanism underlying differential responses of synaptic boutons to lithium is likely to involve distinct modes of SV retrieval. It will be interesting to compare SV cycling and lithium sensitivity in iPSC-derived neurons from both lithium responsive and lithium non-responsive BD patients.

Finally, this study contradicts earlier work reporting no effect of acute and chronic lithium treatment on SV cycling in hippocampal neurons (34). Several reasons could underlie this important discrepancy. First, our measurements are done at 36.5°C rather than room temperature – it is known that the kinetics of SV exocytosis and retrieval are significantly different at physiological temperature (26). Second, Lueke and colleagues use a different pHluorin probe, SynaptopHluorin (SpH). Although SpH and vGpH behave qualitatively similarly, there is a larger fraction of SpH at the plasma membrane in the resting state (35), due in part, to faster diffusion of SpH out of the synapse following SV exocytosis. This could potentially alter measurements of SV cycling, particularly during repeated stimulations. Third, Lueke et al., do not deconvolve SpH transients into SV exocytic and endocytic components using baf treatment, neither do they normalize SpH responses using NH_4_Cl to account for cell-to-cell variation in SpH expression.

In conclusion, our study identifies the SV cycling machinery as a functional target of lithium. Our findings predict reduced glutamatergic neurotransmission in lithium-treated neuron networks, particularly during high frequency bursts of activity, and highlight a novel mechanism by which lithium can mitigate hyperexcitability in BD brains. Our work further pinpoints abnormal SV cycling as a plausible endophenotype of BD, in line with recent studies reporting altered SV dynamics in mouse models of mental illness (19,36). Modelling disease-specific alterations of SV cycling and their reversal by lithium might be one important step toward personalized medicine and drug discovery for mental illness.

## Material and Methods

### DNA constructs, neuron cultures and transfection methods

The pCAGG-vGlut1-pHluorin (vGpH) construct was a gift from R Edwards (UCSF; (18)). Primary rat hippocampal cultures were prepared from E18 embryos according to the Banker protocol (37). These neuron cultures contain ∼ 90% of glutamatergic pyramidal cells and ∼ 10% of gabaergic interneurons. Hippocampal neurons were grown on poly-L-lysine coated glass coverslips (2.5 mm diameter) on top of (but with no direct contact with) a glial feeder layer. Neurons were electroporated with vGpH immediately after dissociation using the Nucleofector kit (Amaxa Biosystems, Lonza) and were imaged 14-16 days after plating.

### Live-cell imaging and field stimulation

Time-lapse confocal imaging was carried out on an inverted Eclipse TE2000-E microscope (Nikon, Japan) with a spinning-disk confocal scan head (CSU-10; Yokogawa, Japan) mounted on the side port of the microscope and equipped with a dual GFP/RFP dichroic mirror. Cells were maintained at 36.5°C on the stage and were imaged the built- in autofocusing system (PFS; Nikon). vGpH was excited using the 488 nm line of an argon ion laser coupled into the scan head by fiber optics. Fluorescence was recorded at 515 nm with a bandpass emission filter (40 nm half bandwidth) and images were captured with an Orca-Flash 4.0 CCD camera (Hamamatsu Photonics, Japan) controlled by MetaMorph 7.8.6 (Molecular Devices, CA, USA) at 0.5 Hz. Samples were imaged using a 60 X (NA 1.4) plan apo objective objective in Tyrode’s buffer (150 mM NaCl, 2.5 mM KCl, 1 mM CaCl2, 1 mM MgCl2, 6 mM glucose, 25 mM HEPES, pH 7.4) supplemented with 25 μM 6-cyano-7-nitroquinoxaline-2,3-dione / CNQX (Tocris Bioscience) and 50 μM D,L-2-amino-5-phosphonovaleric acid / AP5 (Tocris Bioscience). Coverslips were mounted in an RC-21BRFS chamber (Warner Instruments, USA) equipped with platinum wire electrodes. Field stimulation was induced by a square pulse stimulator (Grass Technologies, USA) and monitored by an oscilloscope (TDS210, Tektronix, USA). Trains of action potentials (APs) were generated by applying 20 V pulses (1 ms duration) at 10 Hz. Our typical stimulation paradigm for vGpH measurements involved two consecutive trains of 300 APs at 10 Hz, separated by ∼5 min to allow synapses to recover. vGpH responses between the first and second stimulation were highly reproducible (data not shown). For measuring SV exocytic rates, the second stimulation was preceded (30s earlier) by the addition of 0.5 μM Bafilomycin A1 (Baf; AG scientific, USA). To normalize for total expression of vGpH in each individual bouton, 50 mM NH4Cl (membrane permeable alkalinizing agent) was added at the end of the time series.

### Image analysis

#### SV exo- and endocytic rates

Automated analysis of vGpH transients was previously described in (19). Briefly, responsive boutons were segmented on a ΔF image (F_peak_ - F_baseline_) obtained by subtracting a baseline image (before stimulation) to the image corresponding to the peak vGpH response during the first stimulation (Figure 1b). This differential analysis allowed us to use the same intensity threshold across experiments. We selected a permissive image threshold (0.02%) to ensure segmentation of both strongly and weakly responding boutons with ΔF/F ≥ 5% (Figure 1c). The average vGpH intensity in each segmented bouton was then computed for each time point. Intensity traces were brought down to the same baseline and normalized for vGpH expression by dividing them with the vGpH signal after NH4Cl addition (Figure 1d). The MATLAB script that generates these intensity traces from a single image field is available in supplementary material (Script 1; vGpH_NH4Cl_GRE_v2.m). These intensity traces are then fed to another script (Script 2; exo_endo_fitting_v6_Baf2_GRE.m) that generates the SV exo- and endocytic traces and computes the exo- and endocytic rates, as well as the endocytic time constant τ post-stimulation. This scripts measures a mean vGpH response from all bouton traces in a FOV and overlays the first vGpH transient with the exocytic and endocytic components (Figure 1e). The endocytic component is obtained by subtracting the first vGpH transient from the the vGpH response after Baf addition. The SV exo rate is obtained by linear regression of the first 4 time points (0 - 6 sec) during stimulation (Figure 1f). The SV endo rate is obtained by linear regression of 9 time points (4 - 20 sec) during stimulation (Figure 1e). The time constant *τ* describing vGpH decay after stimulation is obtained by fitting the first vGpH transient to a biexponential function of the form: I(t) = a_1_e^-t/τ1^ + a_2_e^-t/τ2^. τ always refers to the time constant of the fast component. When indicated, vGpH decay was also fit with a monoexponential function.

#### Heatmaps and hierarchical clustering

Heatmaps (Figure 4a) were generated with Python (Jupyter Notebook) using the seaborn data visualization library. vGpH traces from control and lithium-treated neurons were imported as panda dataframes and the sns.heatmap function was used to display these traces as heatmaps with rows corresponding to individual boutons and columns corresponding to time points. Unsupervised hierarchical clustering was performed using the clustermap function from seaborn on a dataset consisting of both control and lithium-treated boutons (Figure 4b).

### Statistical analysis

Population analysis of SV cycling parameters (Figure 2) were measured in 6 to 13 FOVs for each condition (mock-24h, Li-24h, mock-48-72h, Li-48-72h) originating from 4 to 6 independent neuronal preparations. Each FOV typically contained 100-150 individual boutons. More than a 1000 boutons were analyzed for each condition. Cycling parameters for each FOV are listed in Table 1 (Lithium_data_v53.csv, supplementary material). Boxplots were generated with Python (Jupyter notebook) using the seaborn library. The box shows the quartiles (25% and 75%) of the dataset. The whiskers show the rest of the distribution contained within 1.5x the IQR (interquartile range). The median of the distribution is indicated by a horizontal bar. We used the Python SciPy library for statistical analysis of these datasets. The nonparametric Mann-Whitney U test was employed to compare SV rates (exo rates, endo rates and endo time constant post-stimulation) in control and lithium-treated neurons. P values were corrected for multiple hypothesis testing using the post-hoc Bonferroni method. The script (Script 3; Lithium_SV_dataset.ipynb) used to analyze these aggregate data is available in supplementary material).

## Supporting information

Supplemental Script 1

Supplemental Script 2

Supplemental Script 3

## Acknowledgments

We would like to thank Drs Simon Richardson, Giulia Getty, Lauren Pecorino and members of the Fivaz lab for critical reading of the manuscript. This work was funded by research grants to MF from the Leverhulme Trust (RPG-2018-265) and from the Ministry of Education Singapore (MOE2013-T2-1-053).

